# Non-stationary Markovian dynamics shape swim bias fluctuations in zebrafish larvae

**DOI:** 10.64898/2026.06.07.730682

**Authors:** Flutura Shabani, Ashrit Mangalwedhekar, Armin Bahl

## Abstract

Animals are capable of internally modulating behavior as a function of internal state, environmental sensory conditions, and context. Classical models often assume that sensorimotor decisions are driven primarily by external stimuli, with individual biases treated as either constant or evolving monotonically over time. Here, we show that 5-day-old zebrafish larvae display slowly changing directional swim biases even in stable, homogeneous environments, with fluctuations unfolding over many hours. Computational modeling suggests that these biases arise from a non-stationary Markovian process, with two largely independent internal input streams modulating the tendency to repeat swim directions across consecutive swim bouts. These slow fluctuations are present across sensory conditions, including different light intensities and global motion cues, although the switching statistics are modulated by these conditions. Our findings reveal an intrinsic, context-sensitive source of behavioral variability and provide a framework for further study of computational principles that generate spontaneous and adaptive behavior.

## INTRODUCTION

Spontaneous and flexibly adjustable behavior is crucial throughout the development and life of an organism^1–4^. Historically, behavior and its underlying neural basis have been studied in highly controlled and simplified environments, leading to great scientific progress^5,6^. Yet, when interpreting behavioral time series, especially during sensorimotor decision-making tasks, we also need to consider internal states, individual differences, and other modulatory factors^7–10^. Recent technological advances make it possible to tackle the challenges of long-term behavioral tracking and complex analysis, allowing us to study behavior for longer timescales, in unstimulated and naturalistic environments^1,7,11–13^.

Zebrafish larvae are an excellent model to study spontaneous behavior. They swim in bouts, alternating between a brief swim and a pause, at a rate of about once per second^14,15^. A common observation in their trajectories is the tendency to perform several consecutive bouts in the same direction, left or right, before switching, a behavior known as persistence or consecutiveness^16–19^. This tendency has been explored during pharmacological manipulations^20^ and as a function of temperature and illumination intensity^17,21–23^. Previous work has also proposed that consecutiveness can facilitate efficient exploration of the environment, and identified distinct brain regions in the hindbrain that may regulate these transitions^16,24^. Additionally, Markov models were used to describe this behavior, in which left and right consecutive swim states randomly switch with a certain probability^16^. However, it remains unclear whether consecutiveness is stationary over long timescales within individual animals, and whether switching probabilities are symmetrically modulated, or whether more complex statistics are required to describe these slow behavioral fluctuations. Moreover, zebrafish larvae may not always base their swim directions on left or right stimulus information and occasionally switch to a disengaged state, previously captured by a Markov model^25^. Yet, it remains unclear whether and how these state transitions evolve on longer timescales, and how they respond to sensory stimuli in unchanging environments.

Here, we investigate spontaneous behavior in zebrafish larvae at 5 days post-fertilization (dpf) in a uniform environment for continuous periods of 10 hours per individual. We describe this behavior using a Markov model in which switching probabilities are non-stationary. We find that left and right consecutiveness are not symmetric and change dynamically over several hours, suggesting that they are modulated by separate and largely independent internal dynamics. These behavioral dynamics are persistent in different environments, including darkness and global random-dot motion cues. Together, our results open new opportunities to study how neural circuits are modulated over time and how they generate internal-state-driven spontaneous behavior.

## RESULTS

### Swim biases vary considerably over several hours

To explore spontaneous behavior, we recorded freely swimming zebrafish larvae in a uniformly lit environment for 10 hours (**Fig. 1a**). The tracking automatically identified the angular change in orientation during swim bouts, with angles <0° defined as left bouts and angles >0° as right bouts, respectively (**Fig. 1b**). Individuals varied in their overall swim bias, as measured by their probability to turn rightward (*P*(*R*)) over the entire experiment (**Fig. 1c**). There was no population bias. A typical 5-minute-long trajectory shows the tendency of larvae to swim consecutively in the same direction, hereafter referred to as consecutiveness (**Fig. 1d**). Swim bias varied over the 10 hours of the experiment (**Fig. 1e**). To quantify these changes, we analyzed data in 5-min time bins, with a rolling step of 1.6 minutes (5/3) and calculated *P*(*R*) within each bin and fish. We compared results to time-shuffled data of the entire experiment *P*(*R*)_*sh*_ (**Fig. 1f,h** and **Methods**). Across fish, we found that fluctuations in *P*(*R*) were significantly larger and with longer timescales than in *P*(*R*)_*sh*_ (**Fig. 1h–j**), showing that the bias variation did not arise by chance. We repeated analyses with smaller bin sizes and obtained similar, but noisier, estimations (**Fig. S1**).

**Figure 1.**
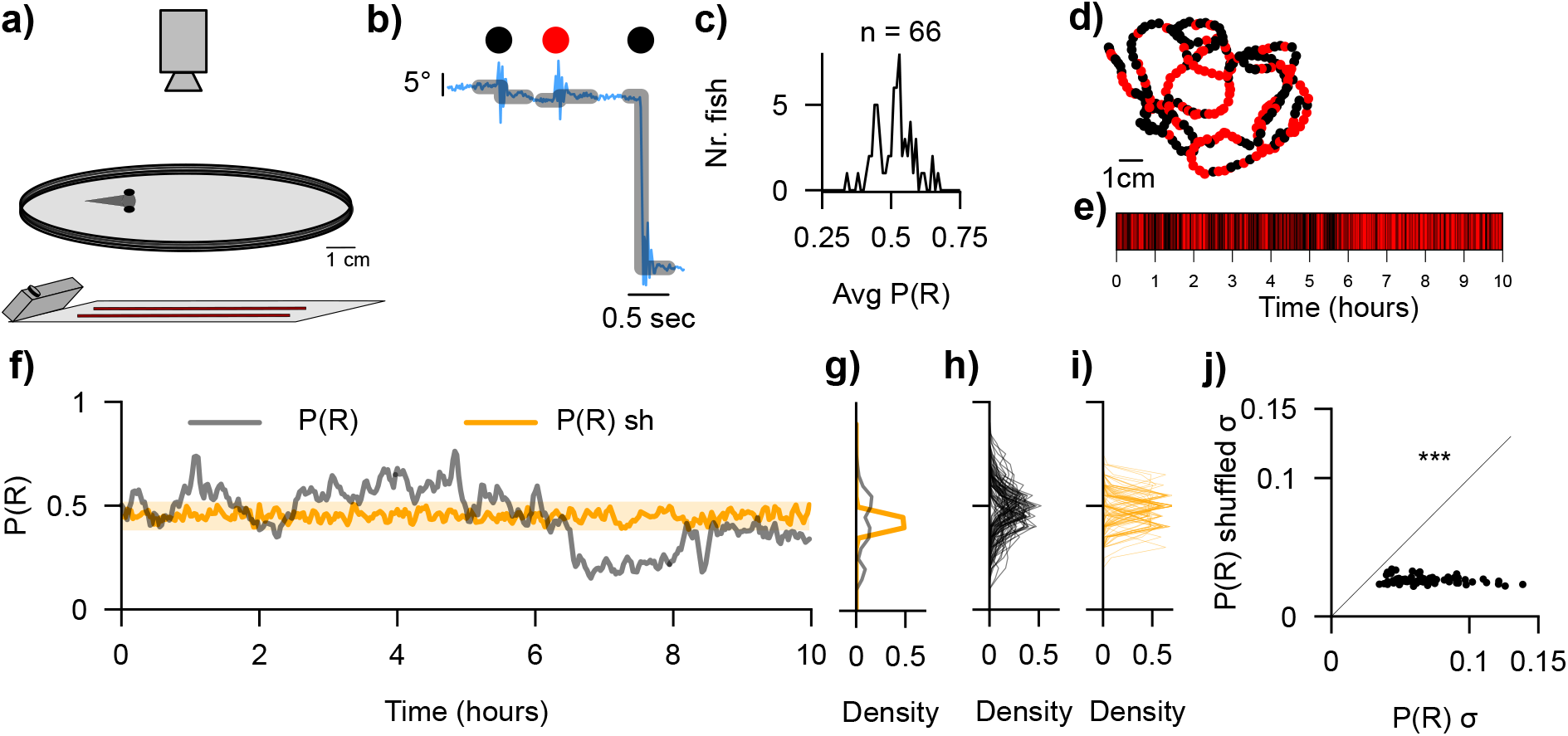
Experimental setup and analysis of long-term swim bias fluctuations. **a**, Schematic of the experimental setup, with the camera recording from above and the projector and infrared lights shining from below. **b**, Definition of right >0° (black) and left <0° (red) swim bouts. **c**, Average *P*(*R*) of each larva in the population (population average 0.506 ± 0.065 std). **d**, 5-minute-long trajectory of an example fish. **e**, Bout directions for a 10-hour-long experiment. **f**, Binned data *P*(*R*) and experiment time-shuffled data (*P*(*R*)_*sh*_). **g**, Probability distribution function (PDF) of *P*(*R*) and *P*(*R*)_*sh*_ of the example fish in (f). **h**,**i**, PDF of *P*(*R*) and *P*(*R*)_*sh*_ of the population. **j**, Population-level standard deviation of *P*(*R*) and *P*(*R*)_*sh*_ (Wilcoxon signed-rank test, W = 2211, p-value: p < 0.001). N = 66 fish in (c,h–j). N=1 fish in (d–g). Also see **Fig. S1**.

Thus, we found that directional swim biases are not stable over time in individual animals but slowly drift over hours, with high heterogeneity across fish (**Fig. 1j**).

### Swim bias fluctuations follow non-stationary Markovian dynamics

We next sought to understand how consecutiveness relates to the dynamics of the observed swim biases in our 10-hour experiments (**Fig. 1**). We quantified the conditional bout-to-bout probabilities *P*(*R*_2_|*R*_1_) and *P*(*L*_2_|*L*_1_) as a measure of right and left consecutiveness, respectively (**Fig. 2a**). Note that this metric quantifies the tendency for 2 consecutive swims and not more. In previous studies, swim bouts were assumed to follow a symmetric Markov model, with temporally stable consecutiveness > 0.5 (**Fig. 2b**)^16^. In theory, however, *P*(*R*_2_|*R*_1_) and *P*(*L*_2_|*L*_1_) may be different from one another and may change over time, even if the fish does not display any bias (*P*(*R*) = 0.5) (**Fig. 2a**). Visual inspection of two example larvae from our dataset showed complex temporal patterns (**Fig. 2c,d,e,f**). In the population, *P*(*R*_2_|*R*_1_) and *P*(*L*_2_|*L*_1_) displayed a wide distribution and a tendency to be >=0.5 (**Fig. 2g,h;** *P*(*R*_2_|*R*_1_): Population average 0.55 ± 0.07 std; *P*(*L*_2_|*L*_1_): Population average 0.54 ± 0.07 std). Since *P*(*R*_2_|*R*_1_) and *P*(*L*_2_|*L*_1_) are symmetric at the population level for all the analyzed variables, to simplify future figure displays, we only show results for *P*(*R*_2_|*R*_1_) unless otherwise specified.

**Figure 2.**
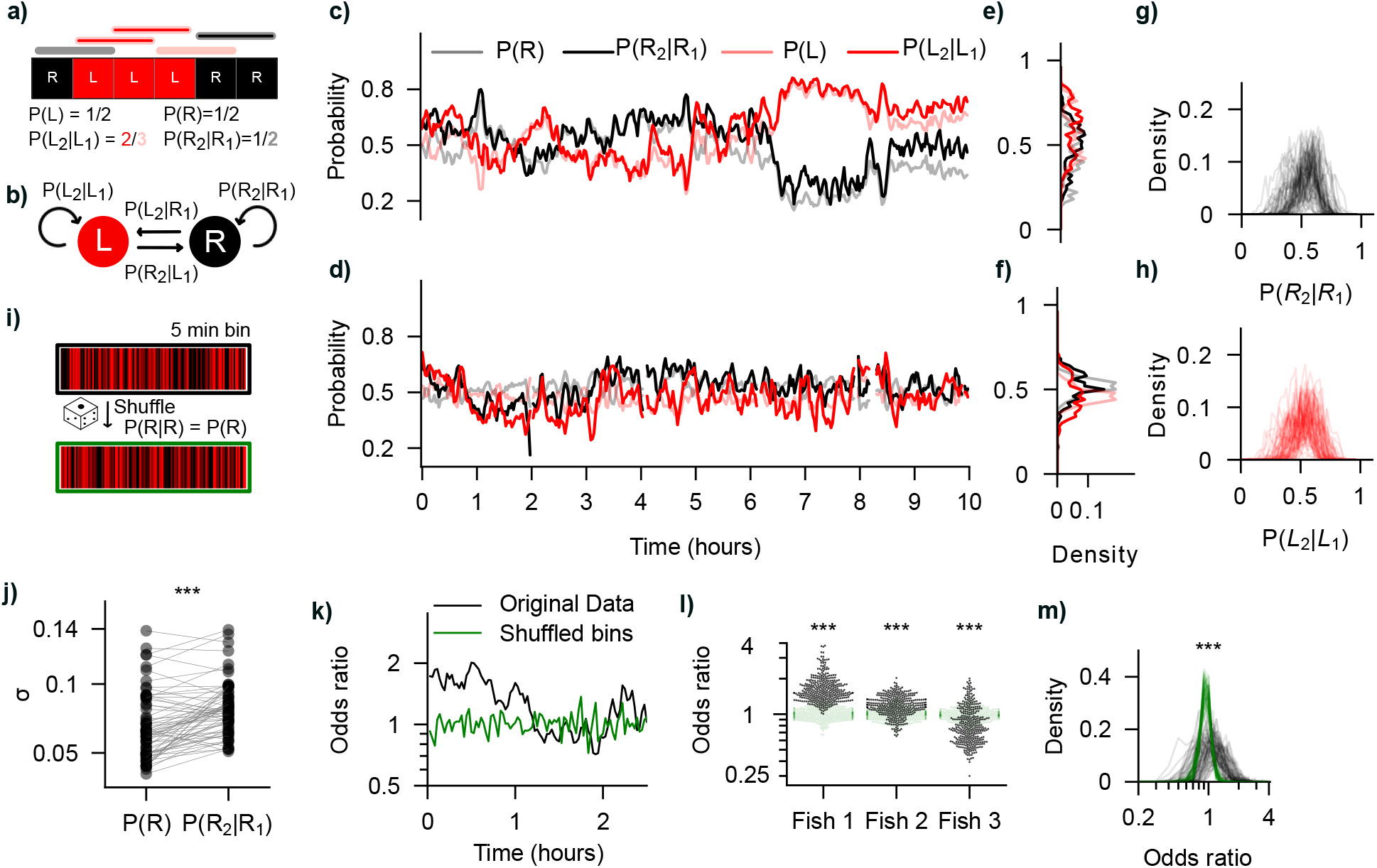
Analysis of non-stationary Markovian dynamics. **a**, Illustration of conditional probability calculation. **b**, Markov model with transition probabilities to remain in a left (L) or right (R) swimming state. **c**,**d**, Biases (*P*(*R*) and *P*(*L*)) and conditional probabilities (*P*(*R*_2_|*R*_1_) and *P*(*L*_2_|*L*_1_)) in 5-min time bins for two example larvae. **e**,**f**, PDF of biases and conditional probabilities for the two example fish in (c,d). **g**,**h**, Both conditional probabilities for each animal in the population. **i**, Illustration of data shuffling within a 5-min time bin. **j**, Standard deviation (σ) of *P*(*R*) and *P*(*R*_2_|*R*_1_) over all time bins for each individual (Population average of σ for *P*(*R*): 0.07 ± 0.02 std; For *P*(*R*_2_|*R*_1_): 0.08 ± 0.02 std; Wilcoxon signed-rank test statistic: 333.0, p-value: p < 0.001). **k**, Odds ratio of *P*(*R*_2_|*R*_1_) relative to *P*(*R*) for an example individual over the first 2 hours of the experiment, together with bin-shuffled data and **l**, for all 5-min bins (individual small dots) for three example fish. (Wilcoxon signed-rank test, statistic: 15, 8480, 16274 for each fish, respectively, and p-value: p < 0.001 for all). **m**, Same as l, PDF for all individuals: Population averages: 1.24 ± 0.19 std and 1.00 ± 0.01 std for original data and bin-shuffled data, respectively. Wilcoxon signed-rank test: W: 2167, p-value: <0.001). N = 1 fish in (c,e,g), another N = 1 fish in (d,f,h). N = 66 fish in (j,m), same fish as in **Fig. 1**.

To better understand how the conditional probabilities change and whether their fluctuations are higher than random measurement noise, we compare the original data to the data with shuffled bouts within each 5-minute bin. Shuffling bouts within a bin preserves the proportion while destroying the ordered pairs of left and right bouts, equating *P* (*R*_2_|*R*_1_) to *P*(*R*) for each bin. (**Fig. 2i**). To show that the fluctuations in *P*(*R*_2_|*R*_1_) are more than an estimation noise artefact from the fish’s general turning bias (measured with *P*(*R*)), we compared their standard deviations across individuals. The standard deviation of *P*(*R*_2_|*R*_1_) exceeded that of *P*(*R*) on average, and particularly in 51 of 66 individuals (**Fig. 2j**), showing that the observed fluctuations are more than estimation noise. Moreover, the lower variability of *P*(*R*) suggests that it could be a consequence of *P*(*R*_2_|*R*_1_) fluctuations (**Fig. 2d** for intuition).

Since *P*(*R*) varies over time, we further used the odds ratio of *P*(*R*_2_|*R*_1_) relative to *P*(*R*) as an additional metric that captures both the magnitude and direction of consecutiveness relative to chance, across the range of values of P(R). In a memory-less process, *P*(*R*_2_|*R*_1_) = *P*(*R*), producing an odds ratio near 1, with minor deviations reflecting estimation noise in a finite sample. Ratios above 1 indicate higher-than-chance consecutiveness, while ratios below 1 indicate lower-than-chance consecutiveness and thus more bout-to-bout alternation of swim direction than expected at chance. Applying this analysis to an example fish over time, alongside bin-shuffled controls **(Fig. 2k**), revealed complex dynamics shaping swim direction, which also varied across individuals in the population (**Fig. 2l,m**). However, since the odds ratio is often <1 (**Fig. 2k,l,m**), the population average does not capture the variability within and across individuals. Beyond this descriptive analysis, *P*(*R*_2_|*R*_1_) and *P*(*R*) are separately parameterized (**Methods**), and, therefore, further comparisons are not meaningful.

In summary, our analyses reveal complex temporal dynamics in directional swimming behavior, showing that Markovian state-switching probabilities are not constant as previously assumed, but fluctuate slowly over time. Next, we quantify the temporal structure of these fluctuations.

### Swim biases and Markovian state transitions fluctuate slowly over time

To describe the temporal structure of the fluctuations in *P*(*R*) and *P*(*R*_2_|*R*_1_), we performed an autocorrelation function (ACF) analysis of both metrics, using the entire experiment-shuffled data for comparison (**Fig. 3a–d**). Autocorrelation decreased over tens of minutes for both, indicating fluctuations of bias and consecutiveness. The number of significant lags was defined as the count of lag indices *k* for which the sample autocorrelation *ρ*(*k*) satisfied 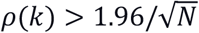, where *N* is the effective sample size. This threshold corresponds to the 95% confidence bound for white noise, based on Bartlett’s asymptotic standard error of 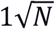 under the null hypothesis of no serial dependence^26^. We found that both *P*(*R*) and *P*(*R*_2_|*R*_1_) have high autocorrelation in time, with high inter-individual variability. Significant lags: *P*(*R*): Avg = 51.46 min ± 23.36 min std, *P*(*R*_2_|*R*_1_): Avg = 54.29 min ± 37.94 min std. *P*(*L*_2_|*L*_1_): 51.72 min ± 39.67 min std, *P*(*R*_2_|*R*_1_)_*sh*_ : Avg = 5.48 min ± 1.23 min std.

**Figure 3.**
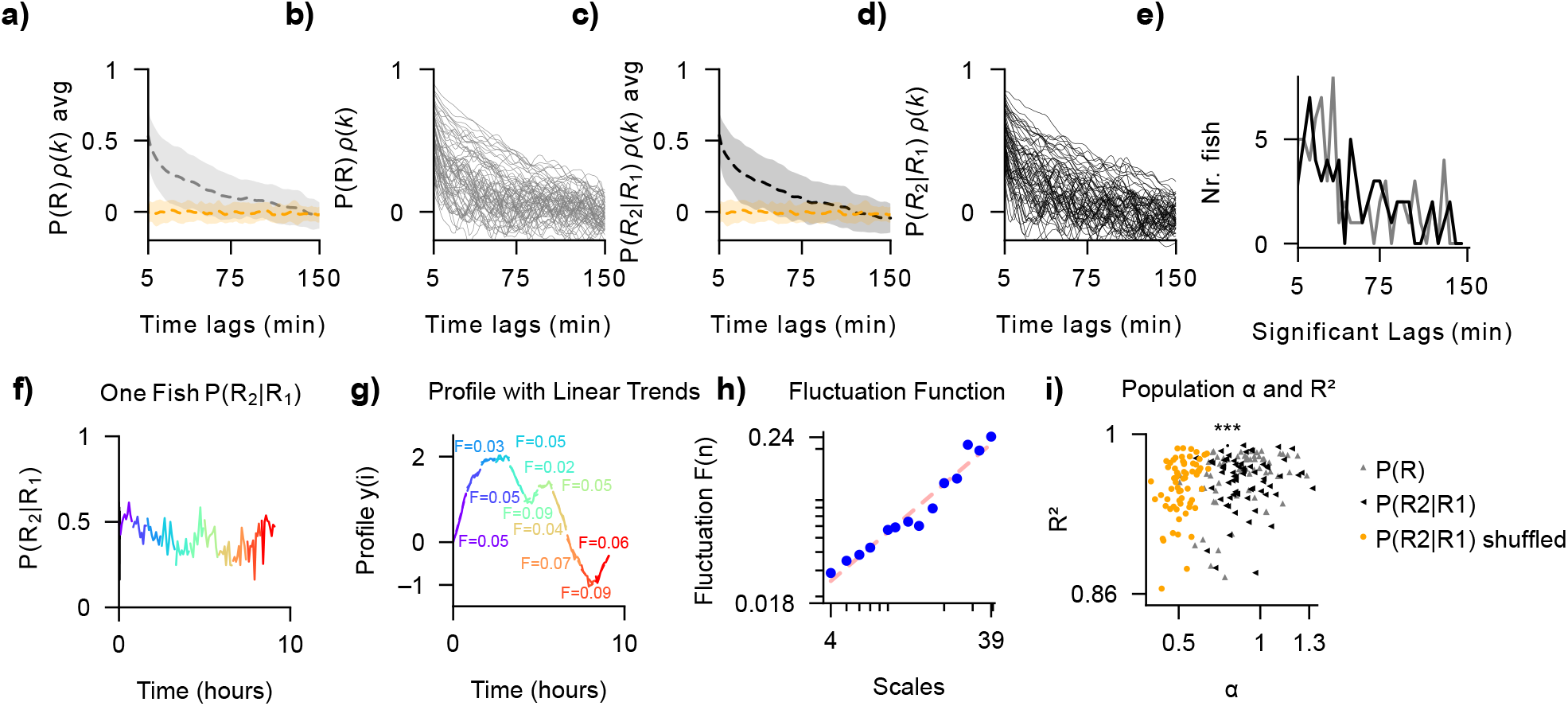
Temporal structure of behavioral biases and Markovian state transition probabilities. **a–e**, Autocorrelation analysis. Colors as in (i). **a**, *P*(*R*) and *P*(*R*)_*sh*_ autocorrelation average and std across fish. **b**, ACF analysis for each individual. **c**,**d**, Same as (a,b) but for *P*(*R*_2_|*R*_1_), respectively. **e**, Significant lags in minutes for *P*(*R*), *P*(*R*_2_|*R*_1_). **f–i**, Detrended Fluctuation Analysis (DFA). **f**, *P*(*R*_2_|*R*_1_) of one example fish with colored segments at scale 10 (10 datapoints per segment). **g**, Profile with linear trends for the example fish in (f). F is the local fluctuation value after linear trend removal for each segment. **h**, fluctuation function *F*(*s*) across the entire experiment at each scale, for the same example fish as in (f), with log-log fitted scaling exponent DFA α = 0.94 (red line). Both axes are in logarithmic scale. See **Methods** for details on DFA and slope interpretation. **i**, The fitted DFA α against goodness of fit (R^2^) for each individual animal. *P*(*R*) and *P*(*R*_2_|*R*_1_) and *P*(*L*_2_|*L*_1_) were compared against shuffled experiment data *P*(*R*_2_|*R*_1_)_*sh*_ using the Wilcoxon signed-rank test and p < 0.001 for all comparisons; p-values were Bonferroni corrected for three independent tests. We did not visualize *P*(*L*_2_|*L*_1_) to simplify the figure display. Data in (a–e,i) from N = 66 fish. N = 1 example fish in (f–h). Same individuals as in **Fig. 1.** Also see **Fig. S2**.

We also used Detrended Fluctuation Analysis (DFA) as a complementary approach to explore scale-dependent temporal correlations in behavioral time series^26,27^ (see **Methods**). DFA quantifies how fluctuations scale with increasing temporal window size after local trends of each window have been removed (**Fig. 3f–h**). In contrast to standard autocorrelation analysis, DFA is relatively robust to non-stationarities in the data. The resulting scaling exponent *α* can then be taken as a proxy for the underlying noise structure. For *P*(*R*), we find that DFA *α* varies largely across individuals with an average *α* = 0.88, ranging from 0.51 to 1.28 (**Fig. 3i**). Similarly, for *P*(*R*_2_|*R*_1_), we found an average DFA *α* of 0.9, ranging from 0.6 to 1.32. For *P*(*L*_2_|*L*_1_), we found an average DFA *α* of 0.9, ranging from 0.62 to 1.24. In contrast, for the shuffled experimental data, *P*(*R*_2_|*R*_1_)_*sh*_ had an average DFA *α* of 0.52, ranging from 0.33 to 0.68. The average *α* values indicate pink-noise-like dynamics, and the shuffled data behave as expected for white noise (see **Methods**). However, there is a large variability in the population. We interpreted that the relatively small sample size per individual^28^, resulting from binning the 10-hour experimental time frame, and the data binning itself^29^ may both have limited our analysis. Yet, the original data *α* was significantly larger than in the shuffled data, suggesting that the behavior could be described with power law noise dynamics, characterized by long-range autocorrelations. We also repeated the DFA analysis for other bin sizes (**Fig. S2**). As expected, estimates of *α* were impacted by the choice of bin size, but in each case, values extracted from experimental data were significantly higher than those from shuffled data.

Hence, both autocorrelation analysis and DFA show that bias and consecutiveness fluctuate over long timescales across tens of minutes. The underlying noise statistics are consistent with power-law noise dynamics, yet our data binning and relatively short experimental duration prevent precise estimation of the scaling exponent. Based on these analyses, we next explored how different variants of Markov models with fluctuating state-transition probabilities may explain our observations.

### State transition dynamics are driven by two internal sources

After showing that the swim biases and internal state transitions are non-stationary with slow fluctuations, we sought to explore the temporal relationship between *P*(*R*_2_|*R*_1_) and *P*(*L*_2_|*L*_1_), specifically, whether their fluctuations over time are independent of one another. To understand their relationship, we generated a set of Markov models with different input configurations, C_1_ and C_2_ (**Fig. 4a–d**, first row). Since DFA showed that the data follows power-law noise dynamics (**Fig. 3h,i)**, we simulated power-law noise with similar fluctuations as *P*(*R*_2_|*R*_1_) and *P*(*L*_2_|*L*_1_) of our dataset (see **Methods**). The detailed explanations of the models are in the **Methods** section. We computed *P*(*R*_2_|*R*_1_) and *P*(*L*_2_|*L*_1_) for each simulated dataset, as we have previously done for our experimental data (**Fig. 4a–d**, second and third row). Next, we compared *P*(*R*_2_|*R*_1_) to *P*(*L*_2_|*L*_1_) between the original data and the models by using the location, spread, and correlation (**Fig. 4a–d**, third row), and used them to manually tune the model simulations, providing us with a quantitative framework to interpret our experimental data.

**Figure 4.**
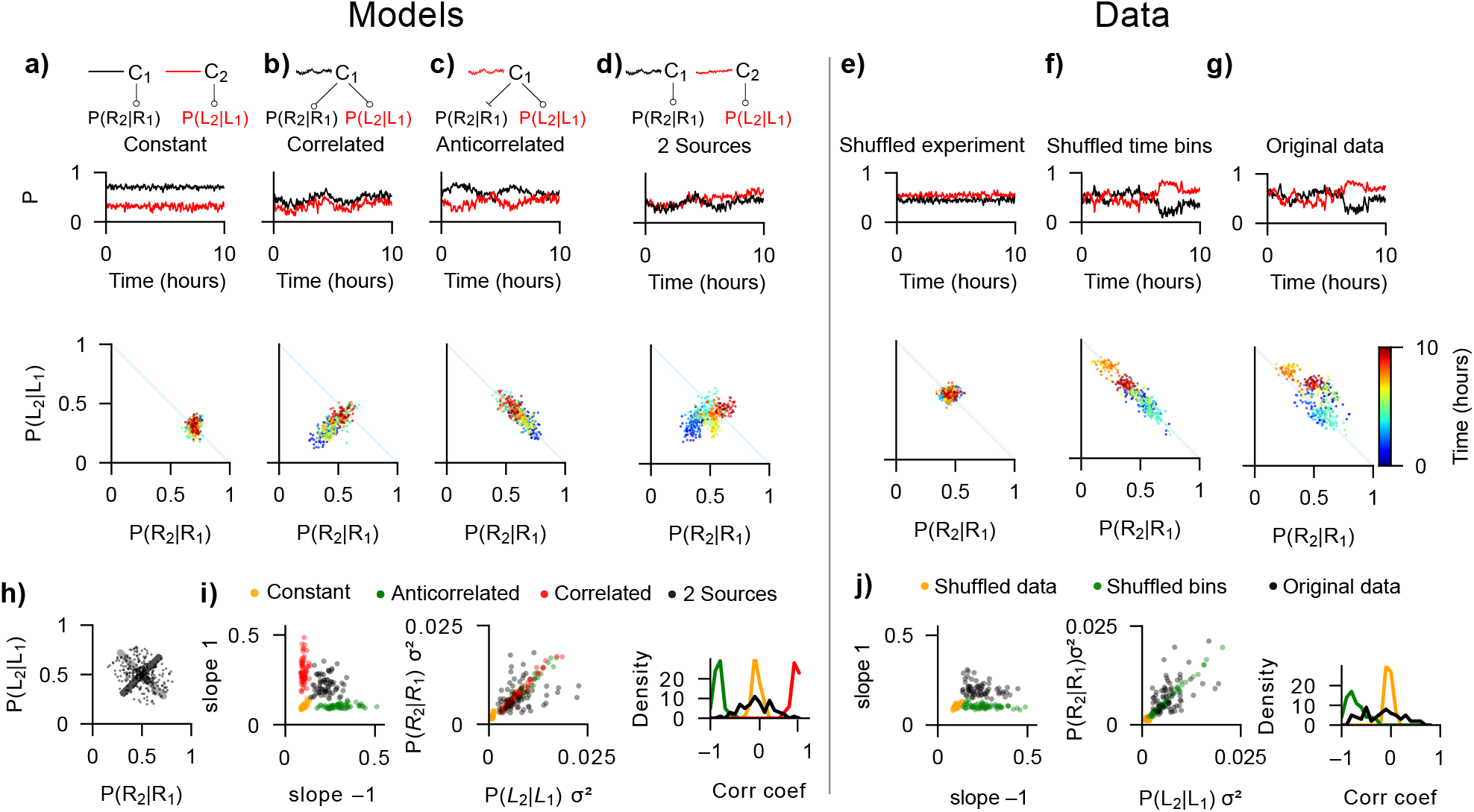
Modeling the relationship between swim transition probabilities dynamics. **a–g**, Individual-level analysis for models and experimental data. First row: Illustration of different Markov model input configurations (a–d). Second row: Calculated swim transition probabilities *P*(*R*_2_|*R*_1_) and *P*(*L*_2_|*L*_1_) fluctuations from individual model simulation (a–d) and from real example fish data (e–g). Third row: Extracted metrics over experimental time for individual example model stimulations (a–d) and real fish (e–g). Simulated and real experimental time is color-coded. **h**, Illustration of how the data range at slopes 1 and –1 is calculated for individual model instances and animals. **i**, Complementary analysis of relationships in the four simulated datasets. Left: Ranges of slopes 1 and –1. Middle: Variances of consecutiveness. Right: Population distribution of Pearson’s correlation of *P*(*R*_2_|*R*_1_) and *P*(*L*_2_|*L*_1_). **j**, Same as (i) but for experimental data. Colors in (i,j) are matched to facilitate the linking of empirical data, shuffling type, and the best-matching model in (i). N = 66 model simulations and experimental fish, respectively, in (i,j). Same fish as in **Fig. 1.** N = 1 example simulation in (a–d,h) and N = 1 fish in (e–g). See **Table S1** for statistical comparisons. Also see **Fig. S4**.

For intuition, we first visually compare the spread of *P*(*R*_2_|*R*_1_) and *P*(*L*_2_|*L*_1_) values in each time bin, in slope = 1 and –1 for one example fish to one example individual simulation for each model configuration. The data of one example fish, when shuffled across the entire experiment, resembled the Constant model, with a very small spread in each direction (**Fig. 4a,e**). In the Correlated model, shown for illustration purposes, the data is primarily spread around slope = 1, and does not resemble the original or shuffled data (**Fig. 4b**). When the data was shuffled only within the 5-minute bins (**Fig. 2i**), it looked similar to the Anticorrelated model, with higher spread in the slope = –1, which would have described our data if *P*(*R*_2_|*R*_1_) was equal to *P*(*R*) (**Fig. 4c,f**). The original data is spread in both directions more than the shuffled data, similar to the 2 Sources model (**Fig. 4d,g**). This showed that neither anticorrelation nor correlation alone can describe the data and that left and right conditional swim bias must be driven by at least two separate fluctuating inputs. To quantify these observations across the population, we use the following three measurements:

First, we calculated the range of *P*(*R*_2_|*R*_1_) and *P*(*L*_2_|*L*_1_) values in slopes 1 and –1 for each individual. If the values are exclusively spread around the slope = 1, they are correlated, and if the values are exclusively spread around slope = –1, they are anticorrelated (**Fig. 4h**). As expected, only the 2 Sources model showed spread in both dimensions, well beyond the chance values based on the shuffled data, because the probabilities were fully independent (**Fig. 4i**, left). The patterns for the Constant, Correlated, and Anticorrelated models were as expected by design. Performing the same analysis on our experimental data revealed that the real fish distribution could be reproduced by the 2 Sources model, in the sense that the population distribution lies beyond the shuffled data equivalents (**Fig. 4j**, left). However, the model does not reproduce the unbalanced ranges in the slope = 1 and –1 that are present in our data, hinting at a small degree of anticorrelation between *P*(*R*_2_|*R*_1_) and *P*(*L*_2_|*L*_1_) in real fish. Fully shuffled experimental data and the data shuffled only within 5-minute bins looked markedly different and resembled the Constant and Anticorrelated models, respectively.

Second, for the Constant, Correlated, and Anticorrelated models, we expected *P*(*R*_2_|*R*_1_) variance to be similar to that of *P*(*L*_2_|*L*_1_) at the individual level, as the temporal fluctuations of inputs are zero or identical in these cases. Only when inputs are independent, as in the 2 Sources model, variances can be different within one individual, which means *P*(*R*_2_|*R*_1_) and *P*(*L*_2_|*L*_1_) do not always show the same degree of fluctuation in one individual, and these expectations matched well (**Fig. 4i**, middle). We then repeated this analysis for experimental data. Again, the 2 Sources model could reproduce the variance distributions of the original data, while the shuffled data matched the Constant and Anticorrelated model configurations (**Fig. 4j**, middle).

Third, Pearson’s correlation distributions between *P*(*R*_2_|*R*_1_) and *P*(*L*_2_|*L*_1_) should also reflect the input structure. As expected, correlations were around zero for the Constant model, and positive and negative for the Correlated and Anticorrelated models, respectively (**Fig. 4i**, right). For the 2 Sources model, the distribution of correlation values was similar to the original data. Again, the experimental data displayed a small shift toward the negative values compared to the 2 Sources model, further revealing the small degree of anticorrelation between *P*(*R*_2_|*R*_1_) and *P*(*L*_2_|*L*_1_). With the limited sample of our 10-hour-long experiment, we cannot fully estimate the true degree of correlation in our data. As an example, in our simulations using the 2 Sources model with fully independent inputs, some individual simulations also produce relatively large correlation values (**Fig. 4i**, right).

To statistically validate our results, we compared the original data to 100 simulations of the 2 Sources model (**Fig. S4**) and computed a Monte Carlo p-value (**Table S1**).

In summary, our analysis suggests that at least two fluctuating sources are needed to drive the dynamics of the probabilities *P*(*R*_2_|*R*_1_) and *P*(*L*_2_|*L*_1_), even though these two sources might not be independent but rather weakly anticorrelated. Having a validated model-based description of how internal dynamics can drive swim bias fluctuations over experimental time, we next explored how environmental cues may shape these features.

### Swim bias fluctuations exist across conditions, yet they are modulated by the environment

So far, we have explored swim bias fluctuations under a dimly lit monotonous environment (**Figs. 1–4**). To assess how robust our observations are across environmental conditions, we next conducted experiments in different brightness settings and with global motion cues (**Fig. 5a**) (see **Methods**). To quantify effects and statistically compare across stimuli, we extracted key behavioral metrics following our previous analysis (**Fig. 5b–h**) and show exact values in **Table S2**. Except for the DFA, which was done with the same data binning as previously (see **Methods**), for the other variables, we used 10-minute bins with 1/3 overlap due to the low swim rate in the Dark dataset. Due to the small sample in individual animals (due to the 10-minute binning), for the DFA of *P*(*R*_2_|*R*_1_), we did not report *α* for individuals in which *R*^2^ < 0.9, for both the original and shuffled data, to increase result reliability. This selection resulted in 5/66 individuals being dropped for the Gray, 53/64 for the Dark, 0/24 for the White, 6/24 for the Noise, and 3/14 for the Coherent Motion conditions, respectively.

**Figure 5.**
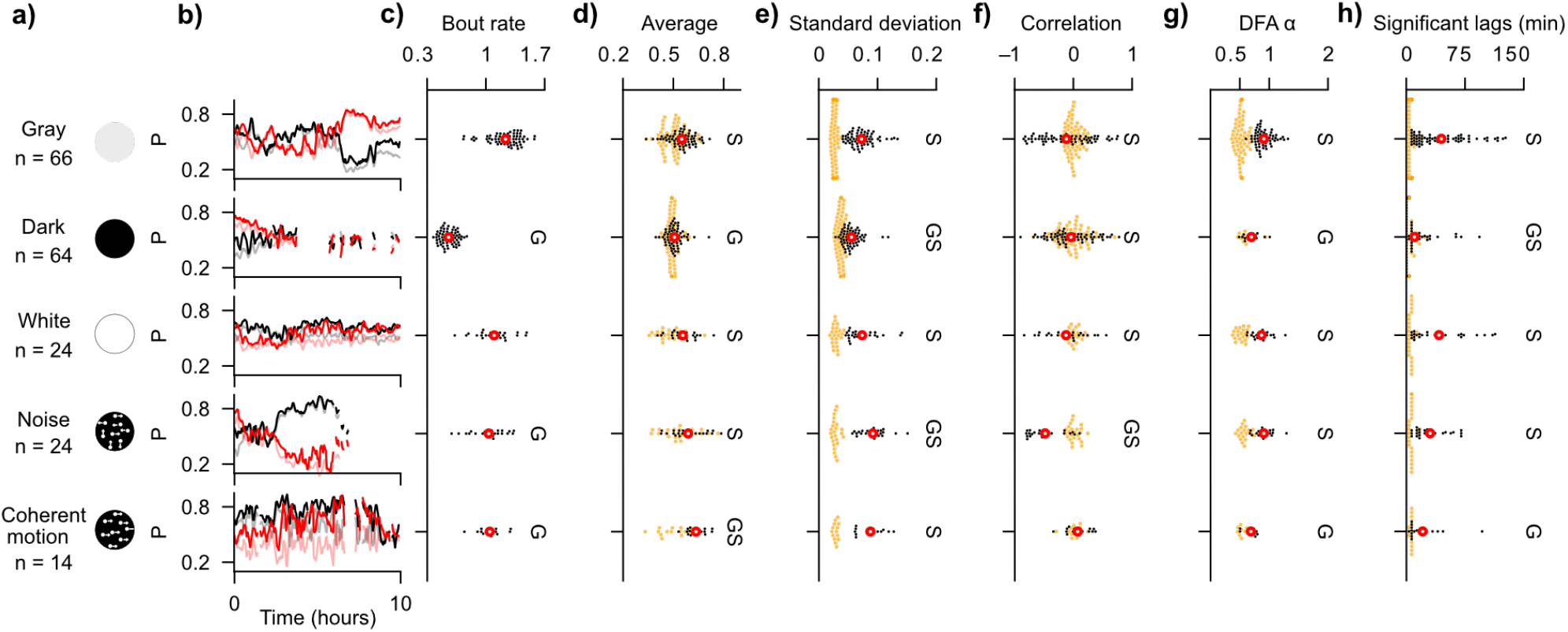
Quantification of behavioral state metrics for different environmental conditions. **a**, Schematic of the experimental stimuli and sample sizes (= number of individuals). Animals in the Gray condition are the same as in **Fig. 1. b**, Extracted probabilities from different example fish over experimental time for *P*(*R*) (pale gray), *P*(*L*) (pale red), *P*(*R*_2_|*R*_1_) (solid black), and *P*(*L*_2_|*L*_1_) (solid red); Same as in **Fig. 2c. c–h**, Population level analysis for each stimulus. Datapoints indicate one individual larva. Original data is shown in black and experiment-wide shuffled data in orange and slightly larger dots. The mean of the original data is shown in red. **c**, Bout rate (average number of bouts per second). **d**,**e**, Average and Standard deviation of *P*(*R*_2_|*R*_1_) over experimental time. **f**, Pearson’s correlation between *P*(*R*_2_|*R*_1_) and *P*(*L*_2_|*L*_1_). **g**, DFA *α* for *P*(*R*_2_|*R*_1_). We only reported *α* for individuals in which *R*^2^ ≥ 0.9. **h**, Autocorrelation significant lags, in minutes, for *P*(*R*_2_|*R*_1_). Significant differences in (c–h) are indicated by letters: G, significant difference compared to the Gray experimental condition; S, significant difference compared to the shuffled data of each experimental condition. For all tests, we performed Bonferroni correction (2 comparisons), such that a value p-value < 0.02 was considered significant, for both G and S. Mann-Whitney U tests are used for all comparisons except for the correlation analysis, in which the Levene test is used. Detailed statistics in **Table S2**. Also see **Figs. S3**, and **S5**.

We first tested behavior in complete darkness (projectors and residual light sources were off, and the room was windowless). Such a configuration also removed the possibility that the visual edge of the arena border may influence the behavioral state transitions we have described (Border effect analysis in **Fig. S3**). The overall swim rate was significantly reduced (**Fig. 5c**), with long periods during which it was too low for reliable estimations, preventing further detailed analysis for these time points (see **Methods**) (**Fig. 5b**, second row). Compared to the Gray condition, the average and standard deviation of *P*(*R*_2_|*R*_1_) were reduced in the Dark (**Fig. 5b,c**). Also, its autocorrelation structure and DFA shifted to lower values, with only autocorrelation being significant in comparison to shuffled data (**Fig. 5g,h**). The correlation values across *P*(*R*_2_|*R*_1_) and *P*(*L*_2_|*L*_1_) were not significantly different from the Gray condition (**Fig. 5f**). These results showed that state switching was also present in darkness, with no visible arena border contrast. However, due to low swim rates, estimation noise might have influenced our analysis, making it seem that darkness weakened fluctuations.

We next probed the behavior with brighter arena light, referred to as the White condition (**Fig. 5b**, third row). Brightness levels have been previously shown to only weakly modulate swim statistics^17,30^, but they may impact bias fluctuations. Effects remained significant in comparison to shuffled data controls, showing that slow swim bias fluctuations also exist under bright light conditions. None of the analyzed variables were significantly different from the Gray condition, even though swim rate was generally lower (**Fig. 5c–h**). Thus, we found that overall light levels had no or very little influence on the internal dynamics of the input streams that regulate the Markovian process.

We also performed an experiment in which we exposed larvae to visual motion noise where the fish swam over a cloud of small dots that continuously drifted in random directions (**Fig. 5a**, 4th row) (see **Methods**). Stimulating the visual input with such constantly changing signals could influence the state transition probabilities and thus override the slow fluctuations in swim bias. Swim rates were significantly lower than in the Gray condition (**Fig. 5c**). Average swim bias was not significantly different from the shuffled data, yet the standard deviation of *P*(*R*_2_|*R*_1_) was higher compared to that in the Gray condition (**Fig. 5e**). Notably, the correlation between *P*(*R*_2_|*R*_1_) and *P*(*L*_2_|*L*_1_) was skewed to negative values compared to shuffled controls (see **Methods**) (**Fig. 5b**, 4th row). This showed that external noise can shift input streams and influence swim statistics in animals in more challenging environments. Other variables were not significantly different, corroborating that swim bias fluctuations are present across environmental conditions.

We next exposed animals to stimuli in which dots coherently moved right-ward or left-ward, relative to the body orientation of the fish, similar to previously used optomotor assays^19,31,32^. Such stimuli are known to reliably induce a tendency to follow motion cues and could thus potentially override spontaneous swim bias fluctuations. We had a relatively small sample of 14 animals. Nevertheless, compared to the Gray condition, we found that the coherent motion significantly increased the consecutiveness, shown by the average *P*(*R*_2_|*R*_1_) (**Fig. 5d**). Additionally, the fish did swim toward the visual motion (**Fig. S5**). This confirmed the above-mentioned previous findings, as the response to strong visual motion increased consecutiveness. If no spontaneous swim bias was present, response to visual motion stimuli that alternate directions every 2.3 minutes, and with an analysis with a 10-minute bin size as in our experiment, could cause small overall fluctuations of *P*(*R*_2_|*R*_1_) around a constant value >0.5, resulting in a small standard deviation. This would be due to the fish swimming toward directional motion cues (left or right). In our results, however, the standard deviation of *P*(*R*_2_|*R*_1_) was not significantly different from the Gray condition, and higher than in shuffled data (**Fig. 5e**), showing that consecutiveness fluctuations were higher than expected if consecutiveness did not change in time. DFA *α* and autocorrelation significant lags were lower than in Gray and not significantly different from shuffled data (**Fig. 5g,h**), suggesting either that the response to the motion stimulus reduced the complexity of the behavior, or it increased the uncertainty of *P*(*R*_2_|*R*_1_) estimations. However, visual inspection showed that the fluctuations were still present at longer time scales than the 2.3-minute motion stimulus lengths and the 10-minute bin size (**Fig. 5b**, last row).

We further explored to what degree the radial position in the arena, time-in-experiment, and swim rate influence behavioral states (**Fig. S3**). In general, fish tended to swim less and be closer to the arena border toward the end of the experiment, shown by the pairwise relationships between time, bout rate, and radial position (**Fig. S3m**,**n**,**o**). Only the radial position was correlated with lower swim bias, meaning that *P*(*R*) got closer to 0.5 when the fish was closer to the border, but not when it swam less frequently (**Fig. S3p,q,r**). A similar relationship was also found for consecutiveness (**Fig. S3s,t,u**). We found heterogeneity in the population in each variable analyzed. Although presence near the arena border weakened bias and consecutiveness fluctuations, these effects were noisy and not sufficient to account for the long-term dynamics, especially considering the presence of the fluctuations in the other experimental conditions, suggesting an intrinsic rather than wall-driven origin.

In summary, by assessing several key behavioral metrics in various environments, we find that swim bias fluctuations persist across different light and motion stimulation regimes. Therefore, we infer that the observed long-term fluctuations are largely driven by internal processes, while they can be externally modulated.

## DISCUSSION

We provide the first evidence that spontaneously swimming zebrafish larvae show a long-term, slowly fluctuating change of swim direction bias over a 10-hour duration in a uniform and non-changing environment. The bias dynamics were captured by a non-stationary Markov model in which left and right switching probabilities fluctuate largely independently over time, following temporally correlated noise patterns. These fluctuations were persistent across environments with different brightness conditions and visual stimulation, while being modulated by sensory stimuli. Our model reproduces key features of the observed swim dynamics, including spontaneous fluctuations in consecutiveness and heterogeneity across individuals.

Previous studies assumed that the Markovian transition probabilities *P*(*R*_2_|*R*_1_) and *P*(*L*_2_|*L*_1_) are mostly stable and symmetric across left and right turns in larvae^16,18,20,33^, changing only in response to temperature fluctuations^21,23^, or asymmetric visual input to the two eyes^17^. We showed that the left and right consecutiveness are neither symmetric nor stable in non-changing environments: they fluctuate spontaneously over long timescales, from minutes to hours, in both uniform (**Fig. 2c,d**) and non-uniform environments (**Fig. 5**). Notably, transition probabilities can also fall below 0.5 (**Fig. 2d**). In this regime, animals display a higher-than-chance tendency to alternate swim directions rather than consecutively repeating them. Such transition probabilities point towards a navigational state in which animals seek to explore long distances rather than staying locally in place^16^. Having flexible transition probabilities should make it possible to switch between local and global exploratory behavior, an idea that can be tested through agent-based model simulations. Between and within-individual variability in spontaneous movement was observed in zebrafish larvae, with variability stabilizing during development as well as in adult zebrafish^34–37^. Our results suggest that some degree of individuality measured in an experiment may be explained by long-term behavioral states, emphasizing the need for longer recordings.

Our 10-hour-long experiment reveals fluctuations over the timescale of hours. Given these dynamics, it is possible that the observed inter-individual variability in our analyses may in fact be a feature of slowly drifting states. Our modelling further suggests that left and right fluctuations are largely independent, though a continuum between correlated and anticorrelated phases cannot be excluded. Additionally, the weak anticorrelation at the population level between the transition probabilities, as seen in real data (**Fig. 4j**, right), is not captured by our model. We cannot fully show if the small anticorrelation is due to limited experimental length or has real biological meaning. Taken together, there is evidence of more complex internal dynamics, which motivates future work with longer automated recordings across days.

Our measured transition probabilities are based on two consecutive swims. Longer sequences can be examined using survival analysis as a function of streak length (the number of preceding bouts in the same direction). Previous work^20^ suggested that the probability of repeating a swim bout direction increases with streak length. We also observe this phenomenon (**Fig. S6**). However, further analyses reveal that such dynamics can also statistically arise when concatenating heterogeneous animal populations with fluctuating behavioral states, and analyzing them as one. Hence, our findings that Markovian dynamics are non-stationary challenge these previous interpretations. Our findings emphasize the importance of quantifying animal behavior from the perspective of the individual and how it changes at smaller timescales.

On the one hand, we used Markov models to describe behavior on relatively short time scales, on the order of tens of seconds, as done previously^16^. On the other hand, our autocorrelation analysis and DFA reveal behavioral state fluctuations at much longer timescales, following noise patterns with 1/f-like temporal correlations. Our DFA results, while limited by sample size and data binning^28,29^, are largely robust across analysis conditions. 1/f-like correlations are observed in many biological systems as signatures of healthy self-organized complexity^38^, including heartbeat and gait dynamics, whose disruption can indicate neurodegenerative disorders^27,39–41^. Thus, statistical analyses of noise fluctuations in larval zebrafish spontaneous behavior can help to further expand the phenotypic perspective in experiments with genetic and drug perturbations^42–44^.

The persistence of swim bias fluctuations observed in the coherent motion experiment (**Fig. 5**), despite the fish following visual motion drift^19,31^ **(Fig. S5)**, suggests that the fluctuations are part of the fish’s core behavioral repertoire, not artifacts of a monotonous environment. Two non-exclusive mechanisms could account for this. First, zebrafish larvae were suggested to alternate between states during which they are attentive or inattentive to visual motion cues^25^, as also found in mice^9^ and humans^8^. They may rely on internal fluctuations for individual decisions during inattentive states, and on external stimuli during attentive states. Second, the motion response could coexist and operate in parallel with internal fluctuations, subtly modulating the direction of each swim bout. Elevated transition probabilities when stimulated with coherent motion are consistent with both mechanisms, encouraging further experiments and modeling to distinguish between them. Importantly, these internal behavioral fluctuations will need to be taken into account when building algorithmic models based on behavioral quantifications. Typically, such model-building has used average responses across many trials, hours, and individuals^19,25,30–32^. However, if subjects flexibly change how they transform sensory cues into motor output over time, averaging-based models will have limited biological predictive power.

### Limitations and future directions

The left-right swim direction discretization used here is a deliberate simplification of the swimming behavior of larval zebrafish. Further analyses may incorporate swim angle, location, bout rate, and bout type^14^. Previous work has identified that the anterior rhombencephalic turning region (ARTR) is involved in controlling turning transitions^16^ and found that activity in this brain region agrees with Markovian dynamics^23^. Moreover, both behavior and neural dynamics are affected by external features, such as temperature^21,24^. Thus, it is likely that our observed bias fluctuations are also represented in the ARTR. Long-term functional imaging over many hours is possible in spontaneously swimming head-restrained larvae, and thus, further work can link non-stationary swim biases to their neural representation.

Human and animal development is characterized by spontaneous and exploratory behaviors thought to be crucial for brain development and motor behavior learning^35,45–47^. It remains to be explored to what degree consecutiveness and bias fluctuations in larval zebrafish behavior generalize across the locomotor repertoire^14^ and how this relates to motor system maturation^48^. Another interesting future direction would be to investigate whether behavioral states of bias persist when larvae begin to feed^7,35^, and how features manifest in adults. Moreover, older zebrafish become increasingly social and tend to stay together^49,50^, which allows them to coordinate movement statistics^51^. It would thus be interesting to explore how individual swim bias fluctuations can potentially influence collective behavior.

This study opens new opportunities to study neural mechanisms underlying internal-state-dependent animal behavior. It is important to describe and model animal behavior and neural dynamics on the level of the individual and, where possible, without temporal averaging. Building such frameworks to have predictive power remains a challenge in neuroscience and behavioral experimentation across species^52^.

## MATERIAL AND METHODS

### Fish rearing and experimental setup

We used wild-type *Danio rerio* (Konstanz strain) larvae at 5 dpf. Larvae were reared in Petri dishes containing E3 solution under a 14/10 hr light/dark cycle at 28 °C. Sex could not be determined at this age. All experiments were conducted from morning to evening, and lasted 10 hours, except for the Noise experimental condition (**Fig. 5a**), in which recordings lasted 8 hours in 16 individuals and 6 hours in 8 individuals. This should be taken into account when interpreting the results. For the experiment, we placed larvae in standard E3 water into custom-designed circular arenas with a 5 mm water level (**Fig. 1a**). The dishes had black walls and a transparent diffusive bottom and were lit from below (AAXA P300 Pico Projector). The room temperature was stably maintained at 26 °C.

The arenas were 12 cm in diameter, with a fish-to-arena diameter ratio of ∼1:24 (fish length ≤0.5 cm). To optimize data storage for our long-term experiments, analysis was performed online, saving only a highly compressed dataset of animal trajectories and swim bouts rather than raw video files. Data was saved, and the experiment was automatically restarted every 2 hours, causing brief tracking interruptions (from a few seconds to a few minutes due to data compression). Within the time frame of our analyses, these missing data are negligible. We had 32 arenas running in parallel, controlled by four computers, with independent stimulation and closed-loop tracking of 32 individual fish at once.

### Behavioral tracking and bout detection

To record the fish, infrared (IR) illumination from below was provided by a 940 nm wavelength LED array. Behavior was recorded at 90 Hz using IR-sensitive cameras (Basler acA2040-90um-NIR) equipped with adjustable macro lenses (Navitar ZOOM 7000) and 850 nm infrared longpass filters (Linghuang Zomei IR 850 nm 52 mm). We used a previously developed tracking algorithm^30^. In short, posture analysis was performed in real-time using custom-written software on Python 3.11 and OpenCV 4.1, using background subtraction to identify the contour. The calculation of head orientation was done by computing the principal component vector of the head for each frame. Orientation data (**Fig. 1b**) were analyzed using a rolling variance (50 ms window). Swim events (bouts) were detected when variance exceeded 1° for at least 20 ms and ended when it dropped below 0.5° for at least 50 ms. Inter-swim interval was defined as the time between consecutive swims (end of bout to next bout start). To exclude potential tracking errors, bouts were discarded if they exceeded the following thresholds: maximum contour area of fish = 600 px, maximum average speed per swim bout = 1.5 cm/s, maximum inter-swim interval = 25 s, maximum absolute orientation change = 150°, and the arena position > 0.99 (normalized radius of arena = 1). Even though zebrafish larvae normally swim around once per second, we permitted inter-swim intervals of up to 25 seconds because fish are sometimes quiescent over the 10-hour-long experiment, in particular in complete darkness (**Fig. 5b**). The bout orientation angles were used to categorize bout direction as right (R) bouts with >0° relative to the body axis and as left (L) when <0°. We analyzed all the data based on this categorization.

### Visual stimuli and experimental conditions

In the Gray (∼600 lux) (**Figs. 1–4**), White (∼1000 lux), and Dark (0 lux) conditions (**Fig. 5**), the arena brightness levels did not change throughout the experiment. Light levels were measured using an LX1330B light meter (Hongkong Thousandshores Ltd., Central Hong Kong). The Dark experimental condition was done in total darkness, with projectors and other residual light sources in the room completely turned off. In the noisy motion and coherent motion conditions (**Fig. 5**), stimuli were dynamically presented throughout the experiment. All visual stimuli were created with Panda3D. Stimuli were shown in random order, either as static or locked to the fish’s position in a closed-loop setup configuration (full closed-loop delay: 60 ms). The random-dot-motion paradigm was used in the noisy and coherent motion conditions and displayed 1200 dots. Each dot was 1 mm in diameter. In the noisy motion condition, the dots drifted constantly in random directions at a speed of 2.6 cm/sec. In the coherent motion condition, stimuli were presented in randomized loops. The loop contained a 20 s baseline period of static dots, followed by a 120 s stimulus, with one of four conditions presented: static dots, coherent lateral motion in relation to the fish’s body axis (leftward, rightward, or forward), or conflicting motion in which dots moved in opposite lateral directions with equal probability; dot speed was 1.8 cm/s in all motion conditions. Condition order was randomized across loops. Except for the left/right motion (**Fig. S5**), the stimuli were not further analyzed.

### Calculation of probabilities

To analyze swim bout direction in time bins (**Fig. 1**), we binned the data using 5-minute bins with a rolling step of 1.66 minutes (1/3 of 5 minutes), resulting in ∼360 bins for each fish for the 10-hour-long experiment, except for the DFA (see below), in which a 5-minute bin size with no overlap was used for all datasets and analyses. The 5-minute bin is used as a trade-off between temporal resolution (**Fig. S1, S2**) and sufficient data in one bin, accounting for the fact that smaller time bins would have more variable sample sizes due to fluctuating swim rates (**Fig. S3a,d**). Additionally, for the comparative analysis between experimental conditions (except the DFA), we used 10-minute bins with a rolling step of 3.3 minutes (1/3 of 10 minutes) to make up for the low swim rate in the Dark condition, while keeping the datasets comparable. The number of bins varied slightly due to the data storage interruption every 2 hours (see above). We chose a bin size and rolling step to have a sufficient number of swim bouts per bin for reliable statistical analysis. The fractions of right swims were calculated for each bin to estimate the probability of a right swim 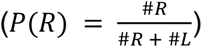. We assigned *nan* values to bins with < 100 bouts (0.33 bouts/second) to increase the robustness of our statistical analysis. Note that *P*(*R*) = 1 − *P*(*L*). Therefore, we did not statistically analyze *P*(*L*).

Conditional probabilities were calculated to assess the tendency for 2 consecutive swims in the same direction (**Fig. 2**). The numbers of each of the 4 possible consecutive pairs (number of left-left (#*LL*), left-right (#*LR*), right-right (#*RR*), and right-left (#*RL*)) were counted. The probability of swimming right given a previous right swim was calculated as 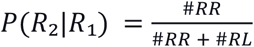, and for the left 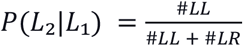, respectively. These estimates are mathematically independent, though finite sample sizes may introduce estimation noise, particularly near boundary values of 0 or 1. For intuition, to calculate *P*(*R*_2_|*R*_1_), we used only pairs that start with R, and for *P*(*L*_2_|*L*_1_), we used only pairs that start with L. *P*(*R*_2_|*R*_1_) is also mathematically independent from *P*(*R*), as the transition probabilities were separately parameterized, as shown by the formulas above. For intuition, apart from the estimation uncertainty on small samples, the way the pairs are ordered tells us nothing about the overall proportion of left and right bouts and vice versa. Importantly, mathematical independence means the parameters are separately defined, and knowing one does not give us information about the other. This does not mean they cannot be empirically associated, which is why we analyze their relationship (**Fig. 4**).

### Data shuffling

To assess statistical significance, the original data were repeatedly compared to the shuffled controls. For full-experiment shuffling, bout directions (R, L) across the entire experiment were randomized using *numpy*.*random*.*shuffle*, after which the same analysis pipeline was applied as for the original data. The number of bouts within each time bin was preserved during shuffling, ensuring that variability in bout rate did not confound comparisons between original and shuffled datasets. To compare *P*(*R*_2_|*R*_1_) to *P*(*R*) (**Fig. 2**), we shuffled bouts within the 5-minute bins. We used this type of shuffling because shuffling the entire experiment generates a constant *P*(*R*) = *P*(*R*_2_|*R*_1_) across bins, making it impossible to statistically quantify how *P*(*R*_2_|*R*_1_) changes relative to *P*(*R*).

### Autocorrelation and Detrended Fluctuation Analysis (DFA)

The autocorrelation function (ACF), implemented in *statsmodels*.*tsa*.*stattools*, was used to calculate autocorrelations for *P*(*R*_2_|*R*_1_) and *P*(*R*) values using 90 lags, as that is when the average of the original data became equal to the average of the shuffled (**Fig. 3a–e**). With data segmented into 5-minute bins and a 1.6-minute sliding window (1/3 of 5 minutes), 90 lags corresponded to 150 minutes. For comparisons between experimental conditions **(Fig. 5h)**, we used 10-minute bins and a 3.3-minute sliding window (1/3 of 10 minutes), and decreased the number of lags accordingly, to 45 lags, which also corresponded to 150 minutes of data. The number of significant lags was defined as the count of lag indices *k* for which the sample autocorrelation *ρ*(*k*) satisfied 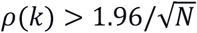, where *N* is the effective sample size. This threshold corresponds to the 95% confidence bound for white noise, based on Bartlett’s asymptotic standard error of 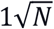 under the null hypothesis of no serial dependence^26^.

We applied DFA to quantify long-range temporal correlations in the time series *P*(*R*), *P*(*R*_2_|*R*_1_) and *P*(*L*_2_|*L*_1_), with a custom-made script based on refs. ^26,27^ (**Fig. 3e–g** and **Fig. 5g**). DFA measures how fluctuations scale across timescales by dividing a signal into windows of increasing sizes, linearly detrending each window, and estimating how residual fluctuations scale with window size (*α*). *α* is indicative of the underlying noise process (see below). All data used for DFA were binned into non-overlapping 5-minute bins, chosen as a middle ground between sufficient data points per bin and adequate temporal resolution, as overlapping bins introduce artificial temporal correlations and inflate the estimated *α* value. The signal was mean-centered and cumulatively integrated, then partitioned into non-overlapping segments across 14 scale levels following a rounded logarithmic progression, ranging from *s* = 4 to *s* = 39 datapoints. Each scale *s* represents the number of data points used for segmenting the data. The segments were locally detrended, and fluctuation magnitudes were computed and averaged across segments for each scale. The scaling exponent *α* was then estimated from a linear fit in log-log space across all scales (**Fig. 3h**). Since at least 2 distinct scales are needed to fit a line in log-log space, and each scale must produce at least 2 segments (⌊*N*/*s*⌋ ≥ 2, where *N* is the number of datapoints), our minimum scale of *s* = 4 and second smallest scale of *s* = 5 together require *N* ≥ 20 datapoints. If at least 2 scales are not possible, *α* is set to *nan* for that individual. To assess the quality of the scaling relationship, the coefficient of determination (R^2^) of the log–log fit was computed.

Regarding the interpretation of DFA *α*: An *α* ≤ 0.5 means the data is not correlated, 0.5 < *α* < 1 indicates that the time series is positively correlated, with *α* ≈ 1 corresponding to 1/*f* noise (pink noise, where the power spectral density is inversely proportional to frequency), and *α* > 1 indicates Brownian-like behavior^26,27^. Individual-level estimates may be unreliable as low sample size and data binning are known to influence DFA scaling exponents^28,29^. They may also explain the discrepancy between our empirical DFA *α* and the model prediction. For the modeling (see below), we use noise generated with a scaling exponent *β* = 1.7 in the frequency domain. This type of noise corresponds to DFA *α* ≈ 1.35 via the approximated relation *β* ≈ 2*α* − 1 ^53^. We did not show direct *β* estimations from our data, as power spectral analysis showed less consistent results than DFA, making it unsuitable for our individual-level estimates.

### Modeling of transition probability dynamics

The aim of our model was to study the relationship between *P*(*L*_2_|*L*_1_) and *P*(*R*_2_|*R*_1_), and to explore if they can be independent. We simulated left/right bout sequences using four Markov models with minute-by-minute varying transition probabilities. For each fish in the Gray experiment, the entire experimental bout data was divided into one-minute non-overlapping time bins, generating *n* bins (∼600 bins for 10 hours), which can vary between fish as previously explained. Each simulated fish drew bout rate and experiment duration from a corresponding real fish, such that the number of bouts at any 1-minute time bin was the same as in the original data. Since our analysis showed that the data follows power law noise, we used it to model the fluctuations of *P*(*R*_2_|*R*_1_) and *P*(*L*_2_|*L*_1_) (**Fig. 3**).

To generate time-varying transition probabilities, power-law noise was generated using the frequency-domain filtering method, in which white noise is transformed into the frequency domain, scaled by *f*^−*β*^, and then inverse-transformed back to the time domain, yielding a signal with spectral exponent *β* = 1.7, reflecting strong long-range temporal correlations close to Brownian noise (*β* = 2)^54^. The value of *β* was manually tuned based on the *P*(*R*_2_|*R*_1_) and *P*(*L*_2_|*L*_1_) scatter plot slope analysis, variance, and correlation values that produce similar dynamics as the real data (**Fig. 4**), and based on visual inspection by checking that the individual model simulations looked similar to the individual fish. The generated power-law noise was normalized to the range 0.2 to 0.8 so that it could be used as probabilities with a comparable range to real data. A subset of *n* consecutive values (number of time bins) was then extracted from a random starting point. Subsetting was done to capture individual variation in the dataset, ensuring that each fish’s probability vectors had meaningfully different effective ranges despite the global normalization. We used a Markov model with four input configuration variants to generate bout sequences, with transition probabilities *f*(*t*) and *g*(*t*) at each time bin. Each configuration produced a simulated dataset, which was then compared across variants. However, the *f* (*t*) and *g*(*t*) vectors were generated once for each individual simulation and then used as input for all of the four configuration variants, in order to keep the model variants comparable. For every one-minute bin (*t*), the number of bouts generated by the models matched that of the corresponding bin in the real data for each individual. The resulting bout sequences were concatenated to produce a single continuous sequence per simulated fish. The model configurations simulated one fish at a time according to the following rules:

1. Constant Model: *P*(*R*_2_|*R*_1_) = 0.3 and *P*(*L*_2_|*L*_1_) = 0.7. We used fixed transition probabilities across all bins.
2. Correlated Model: *P*(*R*_2_|*R*_1_) (*t*) = *f*(*t*) + 0.07 and *P*(*L*_2_|*L*_1_) (*t*) = *f*(*t*) − 0.07. The 0.07 makes sure the values are not exactly equal, though correlated.
3. Anticorrelated Model: *P*(*R*_2_|*R*_1_) (*t*) = *f* (*t*) and *P*(*L*_2_|*L*_1_) (*t*) = 1 − *f*(*t*).
4. 2Sources Model: *P*(*R*_2_|*R*_1_) (*t*) = *f*(*t*) and *P*(*L*_2_|*L*_1_) (*t*) = *g*(*t*).

Simulated bout sequences were analyzed using the same pipeline applied to empirical data, allowing direct comparison of transition probability structures across models.

### Additional notes on data analysis

The odds ratio of *P*(*R*_2_|*R*_1_) relative to *P*(*R*) was calculated for each time bin, as follows:

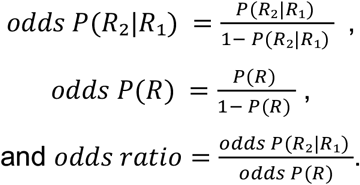

Logarithmic axes were used for odds ratio plots (**Fig. 2 k–m**) to obtain symmetric values around 1, so that equivalent decreases and increases (e.g., odds ratio = 0.5 and odds ratio = 2) are displayed at equal distances.

For the slope analysis **(Fig. 4 i–j)**, the 95^th^ percentile range is used to calculate the range of data points in the slope = 1 and slope = –1 of the scatter plot of *P*(*R*_2_|*R*_1_) and *P*(*L*_2_|*L*_1_). Correlation was calculated with Pearson’s function from the Python scipy package (**Fig. 4 i–j**).

All methods used in previous analyses were also used in **Fig. 5**. For statistical comparisons, each dataset is tested against its own corresponding shuffled data, using a Wilcoxon signed-rank test, and against the Gray dataset, using a Mann-Whitney U test. We additionally used the Levene test (*scipy*.*stats*) for the correlation distributions because most of the datasets were centered at 0, and it was more important to compare the spread of the distributions rather than their means. For the Noise experimental condition dataset, which had a lower average, we used a Mann-Whitney U test.

## Data and code availability

Python scripts for data analyses and simulations, as well as to generate plots, are available at the KonData repository with the persistent identifier https://doi.org/10.48606/eqcq4h1je3mh6vgc. All behavioral tracking data and simulation results generated for this paper can be found at the KonData repository with the persistent identifier https://doi.org/10.48606/f863k8h9pwbvw3hp. Requests for further information and resources should be directed to and will be fulfilled by Armin Bahl.

The above-mentioned persistent identifiers will only work upon final paper acceptance. For the review process, please use the following temporary link containing the data and source code archives:

NextCloud: https://cloud.uni-konstanz.de/index.php/s/gJT3Z7erw5DLkLT

## Acknowledgments

We thank Mohammad Salahsour, Daniel Calovi, and Sophie Aimon for fruitful discussions on data analysis. We thank Daniel Hummel for his useful insight on data analysis and for reviewing the manuscript. We thank Maya Dagher and Sydney Hunt for feedback on the manuscript. We thank Heike Naumann and the animal facility staff for taking care of the zebrafish. We thank members of the Neurobiology and Collective Behavior departments at the University of Konstanz and the Max Planck Institute of Animal Behavior for useful discussions and feedback on this project.

## Author contributions

Conceptualization: F.S., A.B.; Data acquisition: F.S., A.M.; Formal analysis: F.S.; Funding acquisition: A.B.; Investigation: F.S.; Methodology: F.S., A.B.; Project administration: A.B.; Resources: A.B.; Software and data curation: F.S., A.B.; Validation: F.S., A.B.; Visualization: F.S.; Writing: F.S., A.B.; Review: F.S., A.B.

## SUPPLEMENTAL INFORMATION

Document S1

Figures S1 to S6

Tables S1 and S2

**Figure S1.**
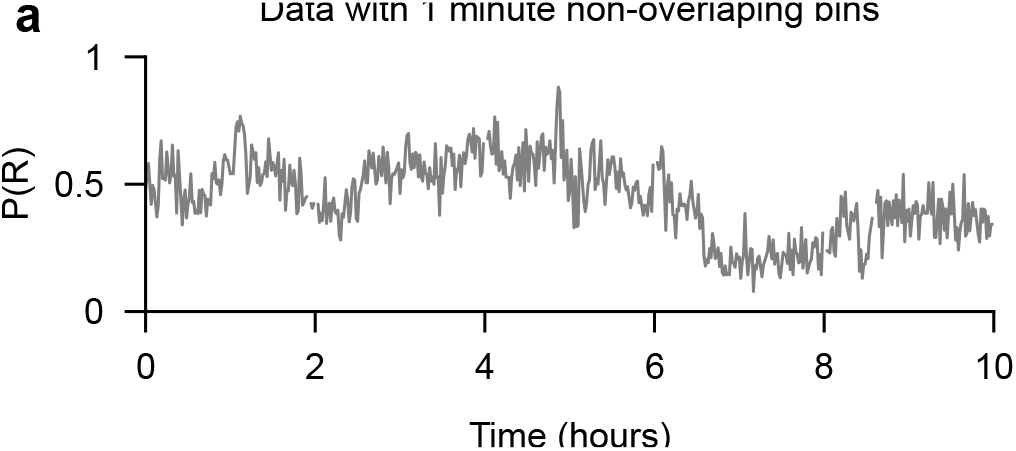
Behavioral analysis with smaller bin sizes. **a**, Segmenting the data from a single example individual (N = 1 fish) into non-overlapping 1-minute bins produced similar, but noisier, *P*(*R*) fluctuations. Same example animal as in **Fig. 1f**. Related to **Fig. 1**.

**Figure S2.**
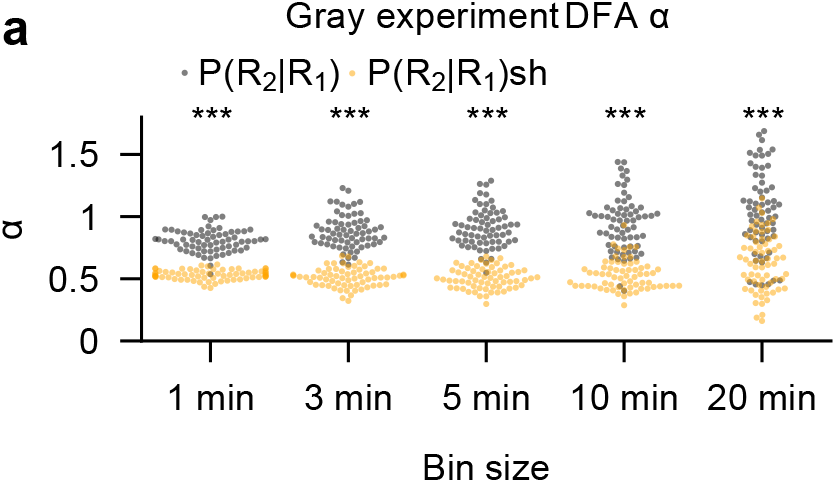
DFA analysis is robust for different bin sizes. **a**, DFA *α* values are shown for individual animals across different non-overlapping bin sizes (timescales) for both the original (black dots) and shuffled data (sh, orange dots). For the original data, the average *α* values for the 1, 3, 5, 10, and 20-minute bins were 0.80, 0.88, 0.91, 0.94, and 1.04, respectively, whereas the corresponding values for the shuffled data were 0.53, 0.51, 0.51, 0.53, and 0.63, respectively. Distributions were compared using the Wilcoxon signed-rank test, with all p values < 0.001 and W = 3, 2, 1, 51, and 187, respectively. N = 66 fish (same fish as in **Figs. 1–3**). Related to **Fig. 3**.

**Figure S3.**
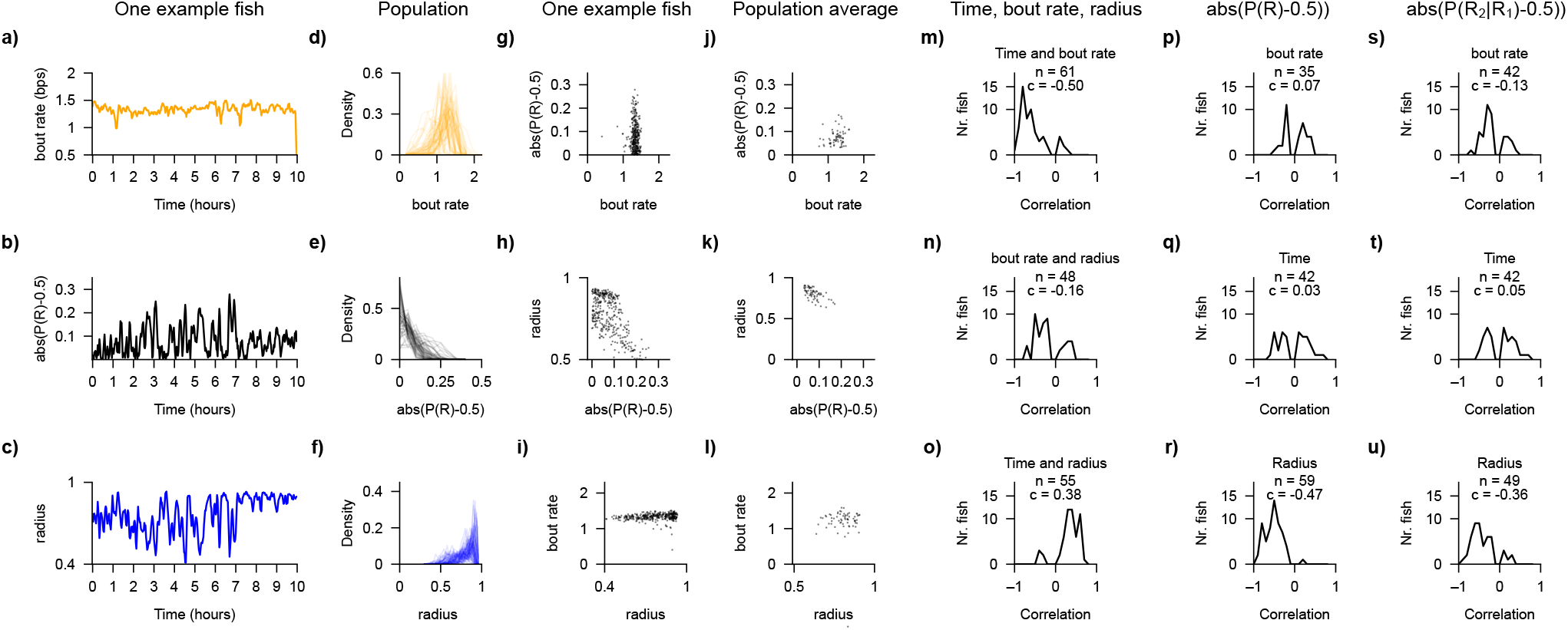
Dependencies between bout rate, arena position, swim bias, and time in the Gray experiment. **a–c, g–i**, One example fish. **d–f, j–l**, population analyses. **a**, Bout rate per second (bps) has low variability in the 10 hours for the example animal. **b**, *ads*(*P*(*R*) − 0.5) is used to measure general bias without considering bias direction. **c**, Radius represents fish radial location in relation to the arena radius, with 0 representing the center of the arena and 1 corner (arena diameter is 12 cm, see **Methods**). **d–f**, Corresponding distributions of each individual in the population for the three variables. **g–i**, One example fish; the relationship between the analyzed variables. Each small dot indicates a 5-minute time bin. **j**,**k**,**l**, Same as in (g–i) but for all animals. Each small dot is a single fish, with values averaged across time. **m**, Most fish swim less in time, as shown by the negative correlation between time and bout rate. **n**, There is a weak negative correlation between bout rate and arena location (radius), indicating that fish swim less near the walls. **o**, Fish tend to be closer to the border of the arena more in later timepoints. **p–u**, The column title shows the common variable on which the correlation analysis is done, and the individual plot titles show the variable unique for each plot. Absolute bias (*ads*(*P*(*R*) − 0.5) is not correlated to bout rate (p) and also not correlated to time in experiment (q), but is negatively correlated to arena location (r). This suggests that biases are more pronounced in the center of the arena. **s–u**, Absolute conditional bias (*ads*(*P*(*R*_2_|*R*_1_)) − 0.5)) is weakly anticorrelated to bout rate, not correlated to time (t), and negatively correlated to arena location (u).

**Figure S4.**
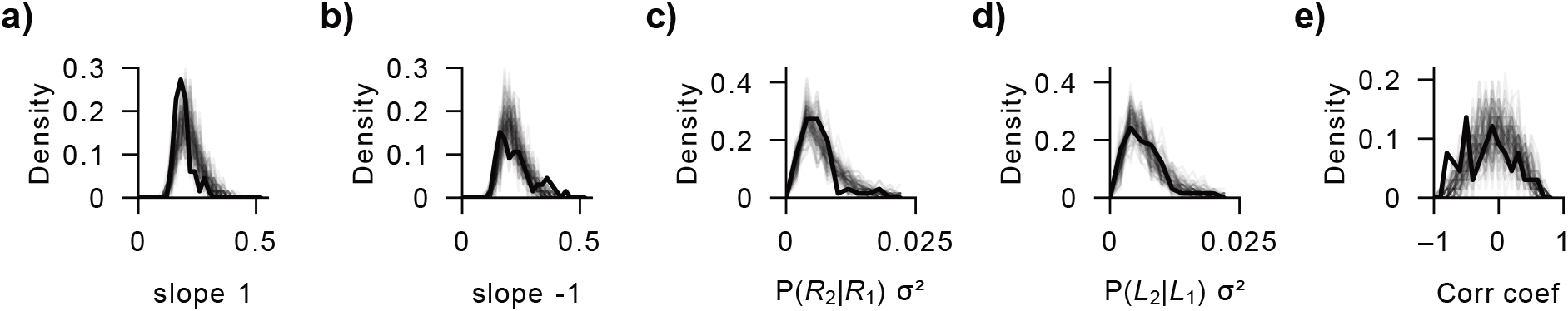
Comparison of original data to 100 simulations using the 2 Sources model. Simulations are shown in semi-transparent gray and original data in black for the variables explained in **Fig. 4. a**, The distributions of the range of values at slope = 1. **b**, The distributions of the range of values at slope = –1. **c**, the distributions of *P*(*R*_2_|*R*_1_) variances. **d**, the distributions of P(L_2_|L_1_) variances. **e**, the distributions of *P*(*R*_2_|*R*_1_) and *P*(*L*_2_|*L*_1_) correlations. Related to **Fig. 4**.

**Figure S5.**
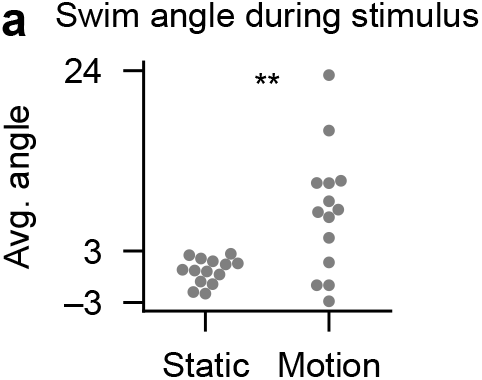
Quantification of the optomotor response. **a**, Average swim angle during left and right stimulus for each individual fish in the Coherent Motion experiment. The average swim angle over 10 hours shows that fish follow directional motion stimulus, although the response is noisy between individuals. Wilcoxon signed-rank test, W=96.000, p=0.002. Related to **Fig. 5**.

**Figure S6.**
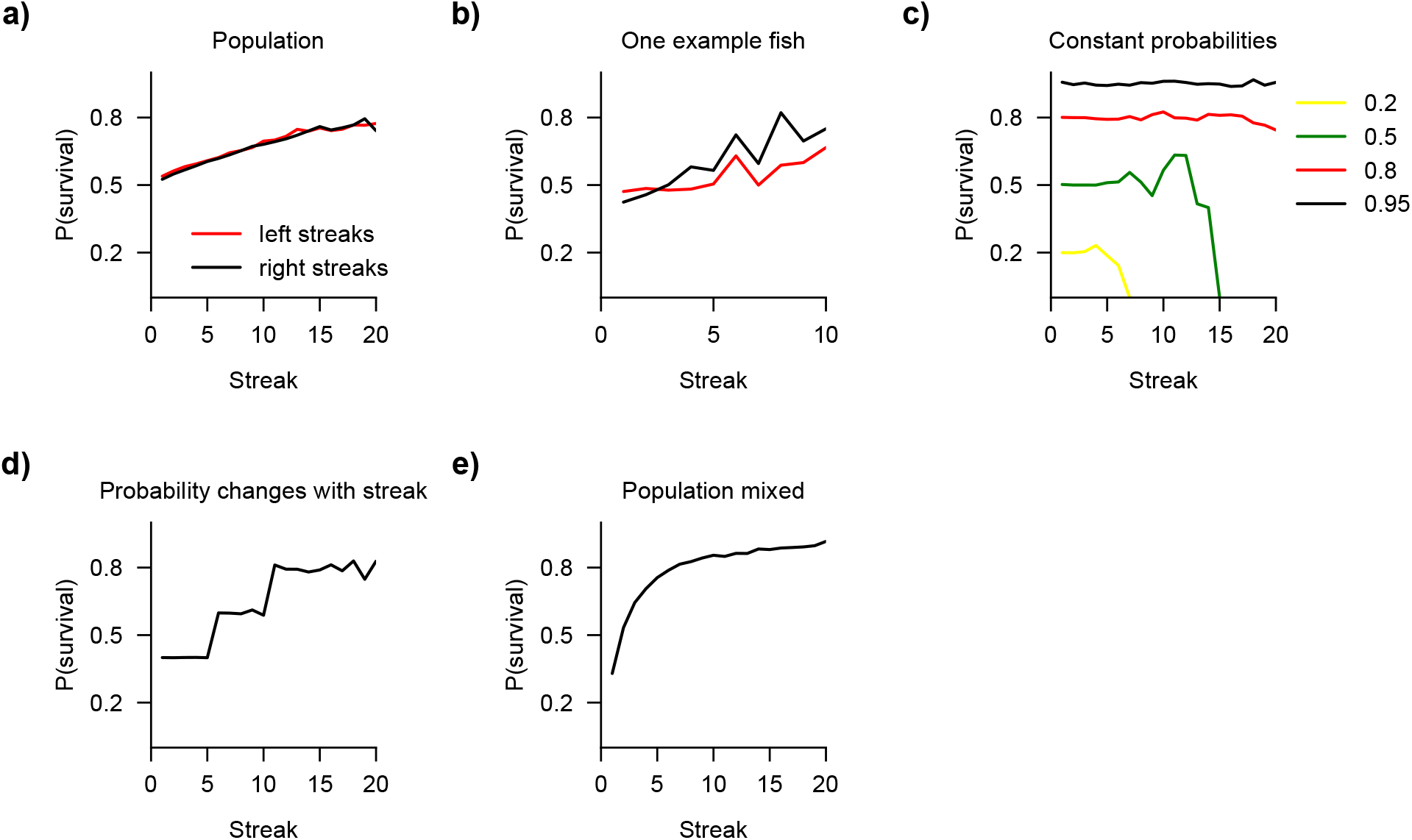
Behavioral state survival analysis of streaks. **a–e**, *P*(*survivial*) represents the probability for a streak of *n* consecutive bouts in the same direction to continue (survive) in the same direction in the next bout (*n* + 1). Real fish data in (a,b) and simulated data in (c–e). **a**,**b**, *P*(*survivial*) calculated from the concatenated dataset of all fish (a) and from one example fish (b). Note that, since survival probability continuously increases with streak length, this analysis gives the impression that the longer the fish swim in the same direction, the more likely they are to continue to do so. As the sample size of streaks in one fish is smaller, the results are noisier, and we stop calculating after streak 10. **c**, We simulated data with a simple coin-flip model with different survival probabilities that are constant over time. Thus, the state survival analysis reveals a straight line. Again, as the sample size gets lower for the longer streaks, the results become noisier. The longest existing streak in a dataset then always has a survival probability of zero. **d**, We also performed simulations, in which *P*(*survivial*) was designed to change with streak length (increasing from 0.4 to 0.6 at a streak of 5, and again to 0.8 at a streak of 10). Analyzing data from this model gives, as expected, the respective streak survival curve. Such a streak-length-dependent transition probability was recently proposed by others (see **Figs. 3B** and **4C** of ref. ^20^ for comparison). **e**, Alternatively, when multiple time series, similar to (c), each with constant probabilities (0.9, 0.6, 0.3, and 0.1) are concatenated and then *P*(*survivial*) is calculated, the same trend appears, as if *P*(*survivial*) changes with streak length. This, however, is not the case for our model. We show that *P*(*R*_2_|*R*_1_), which is the probability of a streak of 2, changes in time in individual animals (**Fig. 2**). Thus, any streak-length survival data needs to be carefully interpreted from that perspective.

**Table S1.** Statistical comparison of the original data to 100 simulations using the 2 Sources model. To reflect the original data distribution, which can have an estimation uncertainty due to the limited number of individuals, we simulated 100 datasets with the same number of individuals as our original data. For each variable, we computed a Monte Carlo p-value defined as the fraction of simulation means deviating from the null mean by at least as much as the real data mean does for each variable. Each simulation mean is the average of a variable across all fish in one simulation run, and the null mean is the mean of the 100 simulation means. The significance threshold was the p-value < 0.02 (highlighted in bold). The real data average is the population average of the same variable in the original fish data. The standard deviation of the p-values for the last 20 iterations is shown (Std. of last 20) as a marker of p-value stability. To summarize, our results showed that the real data exhibits a weak anticorrelation between *P*(*R*_2_|*R*_1_) and *P*(*L*_2_|*L*_1_) which is further shown by the fact that the data is not balanced as the model is, since it has a smaller range at slope = 1 than at slope = –1.

| Statistic | Original data | One simulation | Null avg ± std | Monte Carlo |
| --- | --- | --- | --- | --- |
| **Slope = 1** | Avg: 0.201 ± 0.037 std<br>Range: [0.14, 0.31] | Avg: 0.22 ± 0.05<br>Range: [0.15, 0.35] | 0.224 ± 0.005 | p-value: < **0.001**<br>Std. of last 20: 0 |
| **Slope = -1** | Avg: 0.236 ± 0.073 std<br>Range: [0.13, 0.44] | Avg: 0.23 ± 0.046<br>Range: [0.14, 0.34] | 0.226 ± 0.006 | p-value: 0.12<br>Std. of last 20: 0.038 |
| $P(R_2\vert R_1)$ **Variance** | Avg: 0.0071 ± 0.036 std<br>Range: [0.002, 0.02] | Avg: 0.008 ± 0.004<br>Range: [0.002, 0.023] | 0.007 ± 0.0004 | p-value: 0.15<br>Std. of last 20: 0.022 |
| $P(L_2\vert L_1)$ **Variance** | Avg: 0.0076 ± 0.038 std<br>Range: [0.0026, 0.021] | Avg: 0.0078 ± 0.004<br>Range [0.003, 0.027] | 0.007 ± 0.0004 | p-value: 0.67<br>Std. of last 20: 0.017 |
| **Correlation** | Avg:-0.121 ± 0.381 std<br>Range: [-0.79, 0.65] | Avg: -0.012 ± 0.297<br>Range: [-0.71 0.74] | -0.006 ± 0.038 | p-value: **0.01**<br>Std. of last 20: 0.005 |

**Table S2.**
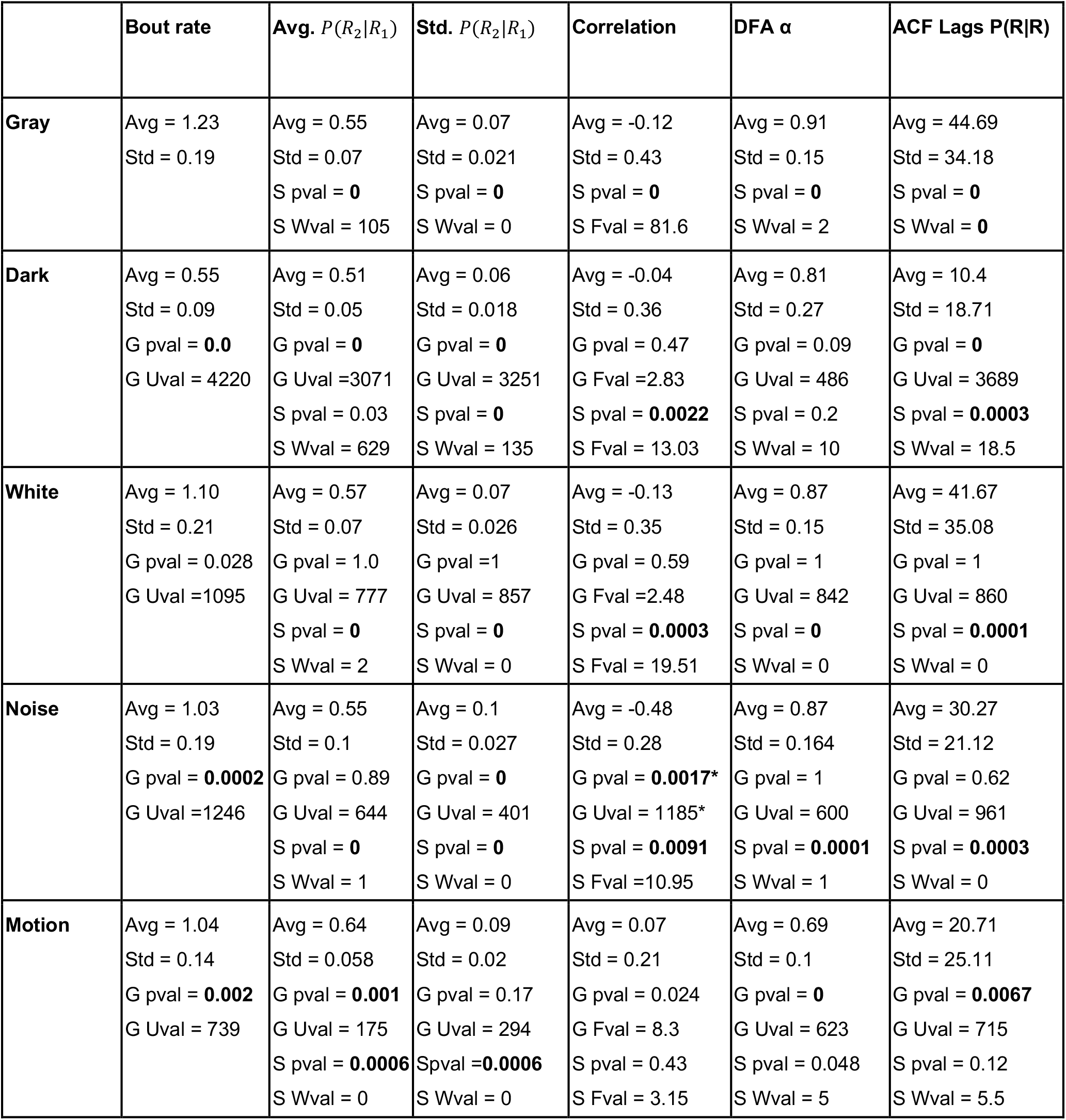
Detailed statistical comparison of values for different experimental conditions related to Fig. 5. Each dataset was compared to corresponding shuffled data (S), using Wilcoxon signed-rank tests, and to the Gray experimental condition (G), using Mann-Whitney U tests. The Levene test was additionally used to compare correlation distributions (see **Methods**). The significance threshold was the p-value < 0.02 (highlighted in bold). Wval - Wilcoxon W value, Uval - Mann-Whitney U value, Fval - Levene test F value. The correlation value for the Noise stimulus, compared to the Gray dataset, the Mann-Whitney p and U value are reported since the average was different (value highlighted with an *).

